# CoDNaS-RNA: a database of Conformational Diversity in the Native State of RNA

**DOI:** 10.1101/2020.10.30.362590

**Authors:** Martín González Buitrón, Ronaldo Romario Tunque Cahui, Emilio García Ríos, Layla Hirsh, María Silvina Fornasari, Gustavo Parisi, Nicolas Palopoli

## Abstract

Conformational changes in RNA native ensembles are central to fulfill many of their biological roles. Systematic knowledge of the extent and possible modulators of this conformational diversity is desirable to better understand the relationship between RNA dynamics and function.

We have developed CoDNaS-RNA as the first database of conformational diversity in RNA molecules. Known RNA structures are retrieved and clustered to identify alternative conformers of each molecule. Pairwise structural comparisons within each cluster allows to measure the variability of the molecule. Additional data on structural features, molecular interactions and functional annotations are provided. CoDNaS-RNA is implemented as a public resource that can be of much interest for computational and bench scientists alike.

**Availability:** CoDNaS-RNA is freely accessible at http://ufq.unq.edu.ar/codnasrna

## Introduction

Many macromolecules alternate between dynamic conformational states to carry out their biological functions. For RNAs in particular, studying this ensemble of conformers is essential to gain the most thorough description of their native state (Ganser *et al*., 2019). The rugged nature of the RNA energy landscape enables functionally relevant conformational changes (Simon and Gehrke, 2009), not only associated with the inherent physicochemical properties of the nucleotide chain, but also triggered by external factors such as changes in temperature or pH and interactions with proteins, metals or small ligands (Mustoe *et al*., 2014). These structural transitions, ranging from local base-pairing perturbations to long-distance domain rearrangements, play a central role in RNA activities such as self-induced folding during transcription, the catalytic activity of ribozymes, the response to altered cellular conditions by riboswitches and the assembly of ribonucleoproteins (Al-Hashimi and Walter, 2008).

In general, structural molecular biology databases do not address the native ensemble of their entries. Some of the many protein structure databases consider conformational diversity explicitly, namely CoDNaS, a collection of alternative tertiary structures of proteins developed in our group (Monzon *et al*., 2016) and PDBFlex (Hrabe *et al*., 2016). While there are databases that compile structural information of RNAs, including valuable resources such as Rfam (Kalvari *et al*., 2018) and URS Database (Baulin *et al*., 2016), these seem to be fewer, less actively maintained or without an easy and comprehensive access to data compared with their protein counterpart. To our knowledge, none of these RNA data resources address conformational diversity explicitly.

In this context, we have developed CoDNaS-RNA to fill a gap in bioinformatics resources to study RNA structure. CoDNaS-RNA constitutes the first RNA database that integrates structural data and annotations to highlight conformational diversity as an essential feature to understand RNA dynamics and function.

### Dataset construction

Each entry in CoDNaS-RNA compiles a cluster of known structures of RNAs with the same sequence, as determined in separate experiments and possibly under different conditions (Figure 1A). Consequently, these conformers can be considered as alternative instances of the RNA structure in its native ensemble. Structural information is obtained directly from mmCIF files in the Protein Data Bank (Berman *et al*., 2000). RNA sequence clusters are built with CD-HIT (at 100% identity and 98% coverage) (Li and Godzik, 2006) and validated with Blastclust (National Center for Biotechnology Information (NCBI). Documentation of the BLASTCLUST-algorithm.) for a robust identification of alternative conformers of the same RNA. All-vs-all pairwise structural comparisons are calculated in each cluster with TMalign (Zhang and Skolnick, 2005). The extent of observed conformational diversity in each RNA is mainly assessed by the maximum RMSD among any given pair of its conformers, with metrics such as TM-score also informed. DSSR (Lu *et al*., 2015) is used to extract all intra- and inter-chain contacts for these conformers, allowing for the comparison of interaction interfaces. External data about ncRNAs is cross-referenced from RNAcentral (The RNAcentral Consortium, 2019) to facilitate further exploration of relevant features. CoDNaS-RNA is first released with 1000 clusters of RNA structures from 2208 out of 3057 validated PDBs (∼77% of available PDB entries), comprising an average of 9.5 conformers per cluster.

**Figure 1.**
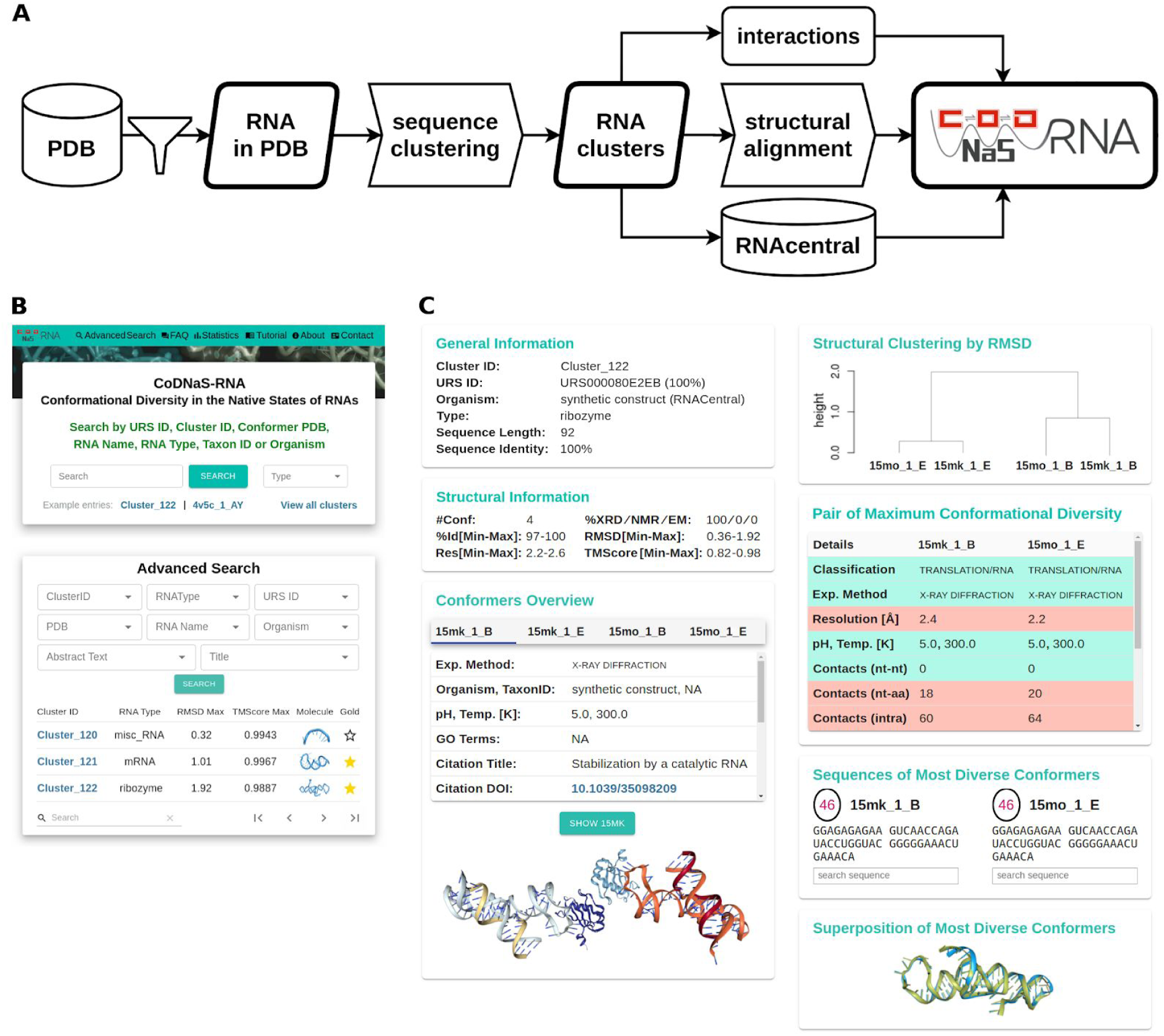
Panel A: Schematic workflow for data acquisition and processing to build CoDNaS-RNA. See details in main text. Panel B: Overview of the basic and advanced search interfaces in CoDNaS-RNA. Panel C: Overview of the main information provided for any selected entry in CoDNaS-RNA. From top to bottom, left column: General information about the selected RNA; structural information about the available RNA conformers and their comparison: and general description of individual confomers. From top to bottom, right column: dendogram of the RMSD-based hierarchical clustering of available conformers; structural comparison of the pair of conformers displaying the maximum conformational diversity as measured by the RMSD between them; searchable sequences of these two confomers; and structural superposition of these two conformers. Panels B and C only show some of the available options and do not necessarily match real entries in the database.

### Usage

CoDNaS-RNA can be searched by one of several criteria (Figure 1B), such as the internal cluster identifier, PDB code, RNA type or name, organism or taxon number, Unique RNA Sequence (URS) identifier, and title or abstract of original publication. Once a cluster is selected from all matching entries, the website provides detailed data on the cluster, organized as separate sections (Figure 1C):

- General information about the cluster e.g. sequence length, cluster sequence identity and source organism. One or more stable URS are assigned to the cluster based on mapping to RNAcentral. The same external resource provides annotations of RNA types.
- Structural information. Provides global statistics from pairwise structural comparisons between members of the cluster. The number of conformers showcases the available evidence on conformational diversity in the cluster. The average, minimum and maximum RMSD and TMscore values, provide the central measurements of conformational diversity.
- Structural overview of conformers, with the full 3D structure for each PDB entry and chain (conformer) in the cluster. These views are presented along with brief structural information and details on the primary citation of the PDB file.
- Dendrogram summarizing the structural clustering of all conformers. Obtained by performing complete linkage hierarchical clustering on pairwise RMSD values.
- Information about the pair of conformers with maximum conformational diversity, i.e. a pair of conformers that display the maximum RMSD value between them. Displays a table of individual and compared data, mainly cross-referenced from the RCSB PDB and taken from DSSR interaction features.
- Interactions on the pair of conformers with maximum conformational diversity. Provides details on the total number of interactions per conformer, presented separately for intra-chain contacts and two inter-chain types of contacts: with other nucleotides and with proteins.
- Sequence information. Shows the sequences of the pair of conformers with maximum RMSD between them, which may differ if at least one is truncated or mutated.
- Interactive view of the structural superposition of conformers with maximum conformational diversity.

The complete datasets used by CoDNaS-RNA can be downloaded as a tab-separated file available from http://ufq.unq.edu.ar/codnasrna/data/codnasrna_v1.0.0.tar.gz

## Conclusions

CoDNaS-RNA is a unique, easy-to-use database to gather information on RNA native ensembles and inspect their conformational diversity. The database showcases possible associations between the diversity in the native ensemble and physicochemical, biological or functional modulators such as origin of the RNA, pH or temperature and binding to ligands. Interaction data is provided at residue level, with at least one conformer in 66% of RNA clusters determined with presence of proteins. CoDNaS-RNA can help to discriminate synthetic conformers from natural ones, while still informing about their shared properties. Moreover, CoDNaS-RNA may also facilitate annotation of RNAs of unspecific types by allowing prospective curators to inspect the annotations of alternative conformers in the cluster of interest. We foresee CoDNaS-RNA can become an essential resource for the wide community of computational and bench scientists.

